# Probabilistic Coverage of the Frontal Aslant Tract in Young Adults: Insights into Individual Variability, Lateralization, and Language Functions

**DOI:** 10.1101/2023.06.03.543491

**Authors:** Wen-Jieh Linn, Jessica Barrios-Martinez, David Fernandes-Cabral, Timothée Jacquesson, Maximiliano Nuñez, Ricardo Gomez, Yury Anania, Juan Fernandez-Miranda, Fang-Cheng Yeh

## Abstract

The frontal aslant tract (FAT) is a crucial neural pathway of language and speech, but little is known about its connectivity and segmentation differences across populations. In this study, we utilized diffusion MRI automatic tractography to investigate the probabilistic coverage of the FAT in a large sample of 1065 young adults. Our primary goal was to reveal individual variability and lateralization of FAT and its structure-function correlations in language processing. Our results showed that the left anterior FAT exhibited the most substantial individual differences, particularly in the superior and middle frontal gyrus, with greater variability in the superior than the inferior region. Furthermore, we found significant left lateralization in FAT, with a greater difference in innervation coverage in the inferior and posterior portions. Additionally, our analysis revealed a significant correlation between the size of left FAT inferior innervation areas and Picture Vocabulary function, highlighting the structural and functional importance of the left FAT in language processing. In comparison, the anisotropy of FAT did not show significant correlation. Overall, our study provides valuable insights into individual and population differences in FAT connectivity and segmentation and sheds light on its critical role in language functions.

## Introduction

The frontal aslant tract (FAT) is described as related to language function recently and has drawn a lot of attentions ^1,2^. Studies have been conducted to identify and reveal the structure of FAT. Catani et al. ^3^ used tractography to first describe the trajectories of FAT. The oblique geometry of FAT accounts for its name *aslant*. Its white matter routes run from the superior frontal gyrus (SFG) at the pre-supplementary (pre-SMA) and supplementary motor area (SMA), curves gradually for about 90 degrees ^4^, and connects the inferior frontal gyrus (IFG), precisely, the *pars opercularis* and *pars triangularis*. The connections between the FAT and the surrounding pathways have been investigated ^5^, such as the arcuate fasciculus (AF), and superior longitudinal fasciculus (SLF). Briggs, Khan et al. ^4^ pointed out that the FAT goes into the AF and SLF complex while they are orthogonal to the FAT in the anterior–posterior plane. The FAT, joint with other associated pathways, facilitates language related and motor planning functions.

Although previous studies have shed light on the structure and function of the FAT, a population-based investigation of *individual differences* in FAT remains to be conducted. While several studies have suggested that the pars opercularis is the primary source of connectivity in the FAT ^3,6^, other studies have shown that in some cases, the pars triangularis may be the primary contributor of fibers ^7^. These discrepancies suggest that there could be substantial individual variation in the distribution of FAT fibers. Given that individual variability is a critical factor in determining the distribution and segmentation of the pathway, it is important to investigate individual differences in FAT. Furthermore, there is a need for a study to explore the relationship between language function and the structural morphology of the FAT. Such a study would require advanced white matter mapping and associated shape analysis on a large cohort.

In this study, we conducted an extensive investigation into the individual variability of the FAT using automatic tractography. Our cohort consisted of 1065 young adults from the Human Connectome Project (HCP), providing a robust sample size for our analyses. To accurately map the FAT bundles, we employed a state-of-the-art automated tractography pipeline ^8^. We took special care to validate the accuracy of our automated fiber tracking pipeline by comparing the results with cadaveric dissection data, ensuring the reliability of our findings. In order to capture the distribution patterns of the FAT within the population, we aggregated the tractography data, allowing us to calculate the population probability of the FAT connections. By quantifying the likelihood of FAT presence and its connectivity patterns, we gained valuable insights into the prevalence and variations of this important neural pathway among individuals.

To delve deeper into the relationship between FAT morphology and language processing, we employed advanced shape analysis techniques ^9^. By applying these techniques to our tractography data, we were able to explore potential structure-function correlations between the shape characteristics of the FAT and performance on two standardized language tests in the NIH toolbox. The first test evaluated reading decoding, namely the Oral Reading Recognition test, whereas the second test evaluated vocabulary comprehension, namely the Picture Vocabulary test. This novel approach allowed us to investigate whether specific structural features of the FAT were associated with variations in language abilities, providing valuable clues about the neural mechanisms underlying language processing.

Our comprehensive study design, combining population-based tractography, validation through cadaveric dissection, and shape analysis, aimed to provide a thorough understanding of the individual variability of the FAT and its potential implications for language processing. By elucidating the intricate relationship between FAT morphology and language abilities, we contribute to the growing body of knowledge on the neuroanatomical basis of language, paving the way for future research and potential clinical applications.

## Results

### Anatomy

We first examined the population-averaged atlas of the FAT ^10^ and its relationship with surrounding pathways. As shown in Fig. 1a, FAT is situated in the frontal lobe, anterior to central sulcus, posterior to anterior ascending ramus (AAR), superior to lateral sulcus and anterior horizontal ramus (AHR). The FAT pathways connect the superior frontal regions, premotor cortex, and pars opercularis. The right FAT has similar geometry and the population-average atlas demonstrates lateralization, which was later examined. The FAT is depicted in orange in the white matter cortex specimen in Fig. 1b, innervating pars opercularis (yellow), pars triangularis (green), and premotor cortex (white). In Fig. 1c, an anatomy dissection image from another subject demonstrates the relationship of FAT and its surrounding white matter fiber pathways. FAT is positioned more medially to the SLF II (green), SLF III (red) and AF (yellow), while the SLF I (pink) is medial to the FAT. The SLF I, SLF II, and SLF III runs in the orthogonal direction to the FAT. The anterior part of the AF also runs in the orthogonal direction to the FAT, but its posterior part has a curve of approximately 90 degrees and becomes parallel to the orientation of the FAT.

**Figure 1.**
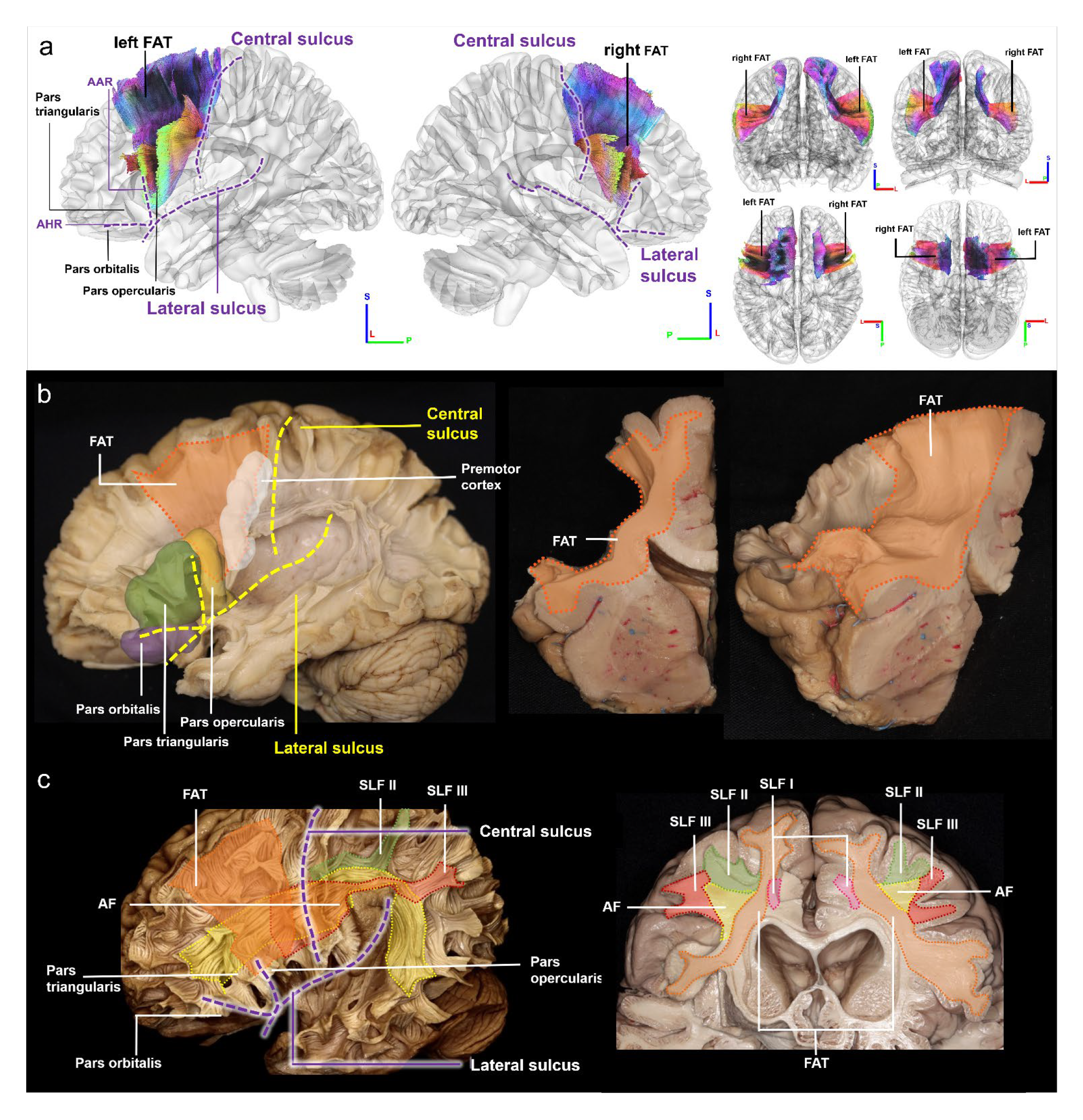
Population-averaged FAT and its spatial relations to cortical regions, sulci, and surrounding white matter fiber pathways. (a) The location of FAT is in the frontal lobe, anterior to central sulcus, posterior to anterior ascending ramus (AAR), superior to lateral sulcus, anterior horizontal ramus (AHR), and pars orbitalis, innervating the premotor cortex and pars opercularis. (b) FAT can be visualized in the white matter cortex specimen (medial frontal gyrus has been removed to facilitate visualization). The coronal view of the FAT shows the white matter fibers running from the SFG to the IFG. Lateral view of the FAT visualizes the fiber tract connecting the pre-supplementary and supplementary motor cortex areas to the inferior frontal gyrus. (c) FAT and other tracts are color coded and visualized in the white matter cortex specimen. The location of FAT runs more medial to other surrounding structures, such as SLF II, SLF III and AF, whereas SLF I is medial to the FAT.

### Population probabilities

We conducted a probabilistic analysis of individual differences in the coverage of the FAT among 1065 young adults. Fig. 2 displays the population probabilities of FAT coverage at the 1st, 2nd, and 3rd quantiles (Q1, Q2, Q3), where darker colors indicate higher coverage percentages. Our findings reveal a significant variability in FAT coverage, particularly in the superior frontal and middle frontal gyrus, with larger differences observed in the superior portion. Additionally, we observed relatively minor variations in the FAT regions proximal to the central sulcus. We quantified the volume of Q3, Q2, and Q1, with volumes of 8932 m^3^, 19608 m^3^, and 32740 m^3^, respectively. Q3 represents FAT coverage that were shared between more than 75% of the population, whereas Q2 and Q1 represent those shared between 50% and 25% population, respectively. Q2 has 119.53% more coverage volume than Q3, while Q1 has 66.97% more coverage volume than Q2, suggesting that there are substantial individual differences, mostly at the anterior portion of the FAT.

**Figure 2.**
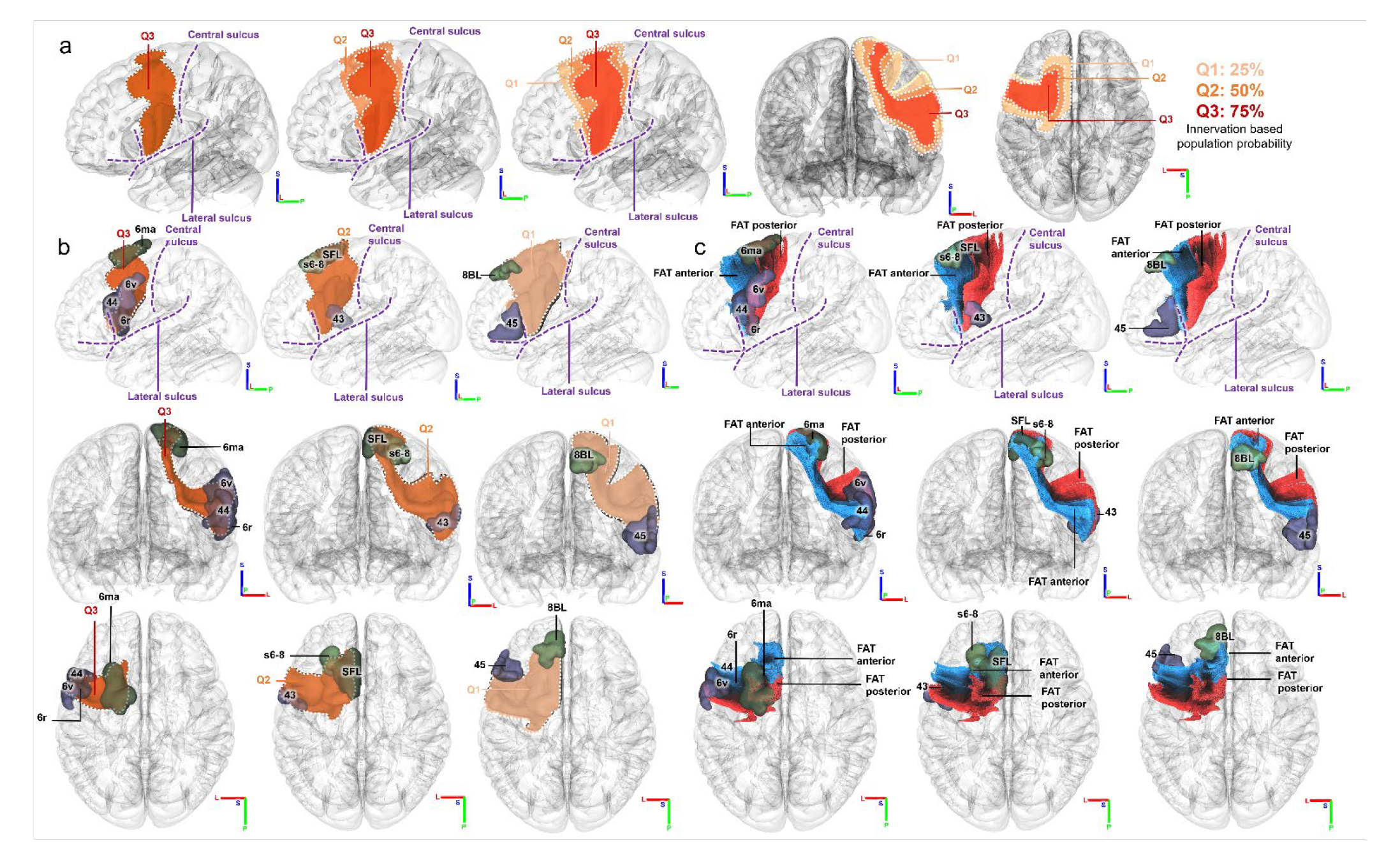
Population probabilities of FAT coverage at 1st, 2nd, and 3rd quantiles (Q1, Q2, Q3) of the human young adults. (a) Q2 has 119.53% more coverage in volume than Q3, whereas Q1 has 66.97% more coverage volume than Q2. There are substantial individual differences in FAT coverage in the young adult population, particularly at the anterior portion of FAT. (b) Q3 mainly innervates area 6ma, 44, 6r, and 6v in HCP-MMP atlas. Besides the areas that Q1 innervates, Q2 also innervates s6-8, SFL, and 43. In addition to the areas that Q3 and Q2 innervates, Q1 innervates 8BL and 45 as well. (c) Among all the areas Q3 innervates, 6ma innervates both posterior and anterior FAT, while 6r and 6v innervates the posterior FAT, 44 innervates the anterior FAT. For the Q2 innervated areas, SFL innervates both posterior and anterior FAT, while 43 innervates the posterior FAT, s6-8 innervates the anterior FAT. Within the areas Q1 mainly innervates, both 8BL and 45 innervates the anterior FAT.

We further used the HCP-MMP atlas to identify cortical regions. As shown in Fig. 2b and 2c, our results show that Q3 innervates regions 6ma, 44, 6r, and 6v. This suggests that the majority of population has these regions innervated by FAT. We roughly segment the FAT into anterior and posterior parts based on the innervated areas. Both parts innervate 6ma, whereas only the posterior FAT innervates 6r and 6v, and only the anterior FAT innervates 44. Furthermore, Q2 innervates s6-8, SFL, and 43, with SFL innervating both posterior and anterior FAT, while 43 innervates only the posterior FAT, and s6-8 innervates only the anterior FAT. Additionally, Q1 innervates 8BL and 45, in addition to the regions innervated by Q3 and Q2. Both regions are innervated by the anterior FAT, suggesting that most individual differences are located at anterior FAT around 45 and 8BL.

Fig. 3a illustrates the lateralization of the frontal aslant tract (FAT) coverage based on population probability. The first two rows demonstrate that the left FAT has a larger volume in all three quantiles, while the third row presents a mirrored image of the left FAT on the right side for ease of comparison. Our findings indicate that the left FAT exhibits a greater degree of individual variability than its right counterpart. Furthermore, more individual differences have emerged in the inferior and posterior innervation area, highlighting significant disparities between the left and right FAT.

**Figure 3.**
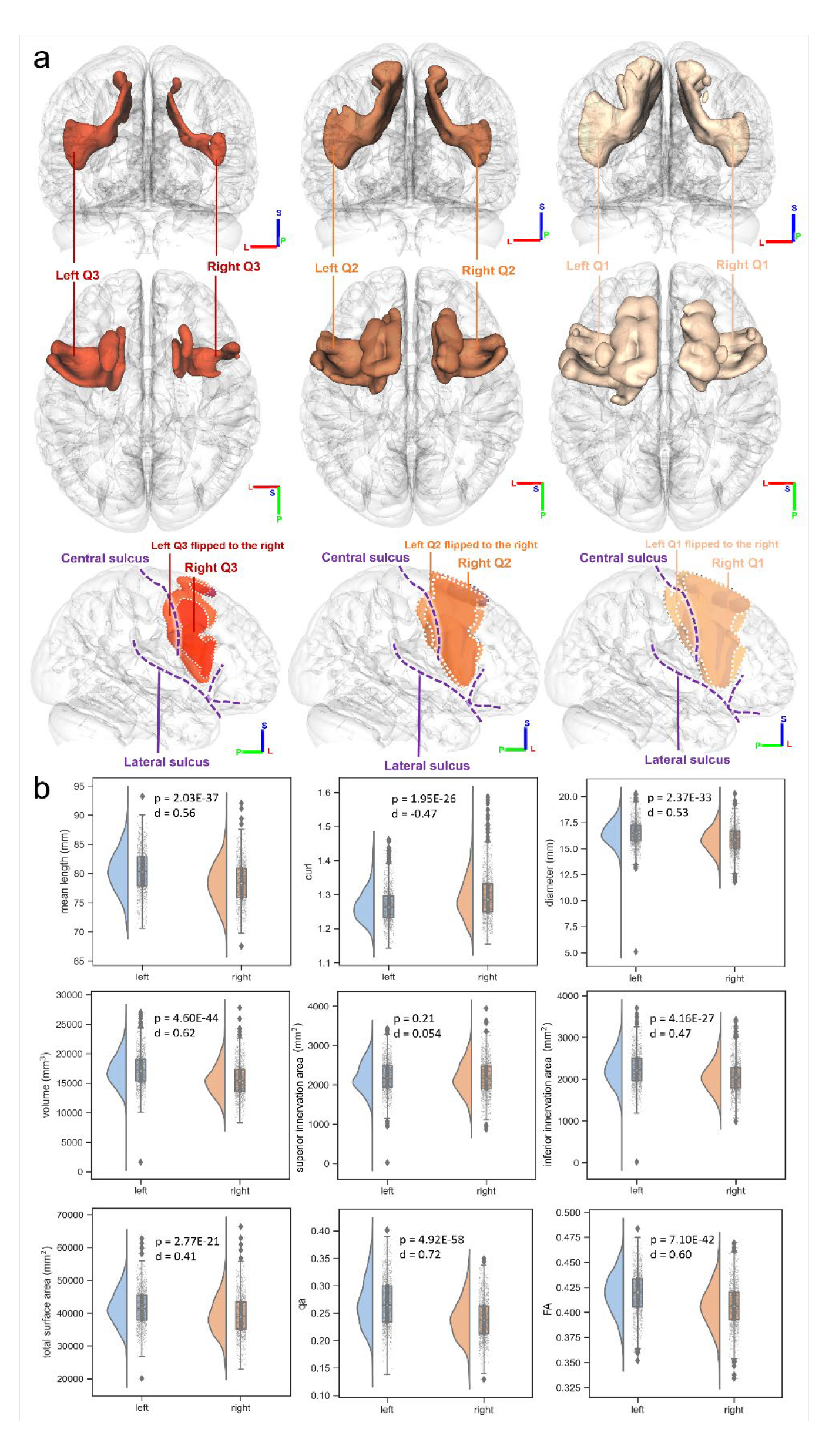
Comparison between the left and right FAT. (a) The population probabilities of FAT coverage shows that the left FAT has a larger individual difference compared with the right FAT. The first two rows show that the left FAT has a larger volume in all three quantiles, while the third row mirrors the left FAT to the right side to facilitate comparison. The results shows that the left side exhibits a greater degree of individual variability. More prominent disparities in individual differences between the left and right FAT arise in the inferior and posterior innervation area. (b) The statistics comparisons of the left and right FAT and their Cohen’s d showing lateralization. Although most of the covariates are left biased, the curl of the FAT is right biased. The superior innervation area has the least lateralization, while the QA of the FAT has the most lateralization.

Fig. 3b further compares shape and diffusion metrics between the left and right FAT in 1,065 young adults. The superior innervation area did not display a significant difference (p=0.21), along with a small effect size (d=0.054), while all remaining metrics showed strong significant differences (p<0.001) with medium effect sizes (Cohen’s d ranged from 0.4 to 0.7). All metrics demonstrated left dominance, with the exception of curl, which exhibited a more curved structure in the right FAT. Notably, there were distinct differences in the inferior innervated areas, consistent with the findings in Fig. 3a. Among all the statistics, QA exhibited the most significant difference with a Cohen’s d value of 0.72, suggesting that the left FAT is a more compact fiber bundle. Overall, FAT exhibits significant left lateralization in both macroscopic shape measures and microscopic diffusion metrics.

We presented the population probabilities of the FAT’s innervation with various cortical regions and specified its coverage area in Fig. 4. These probabilities were calculated based on a tract-to-region connectome using the HCP-MMP parcellations ^8^. The cortical regions were color-coded to indicate different innervation compartments of the FAT bundles and regions. Specifically, green represented superior innervation, blue represented inferior innervation, and purple represented frontoparietal innervation. Fig. 4b, 4c, and 4d showed brain areas of the left hemisphere, and the probability of innervation for each area was quantified. Area 44 had a higher probability of innervation in the inferior region over area 45, but both had a high population probability of over 90% in innervation with the FAT, indicating that most people have FAT connections in both areas. Fig. 4e showed the population probability difference of the FAT coverage between left and right, with the circle’s surface area proportional to the average volume of the area. The colors are coded in the same way. A majority of the regions, thirty-one in total, were left-biased in percentage, while the most biased ones including areas FEF, 45, 55b, and 4. These areas are located in the anterior and posterior parts of the FAT, further supporting the conclusion drawn in Fig. 3 that there is a greater degree of individual variability in this region, thereby underscoring the notable disparities between the left and right FAT.

**Figure 4.**
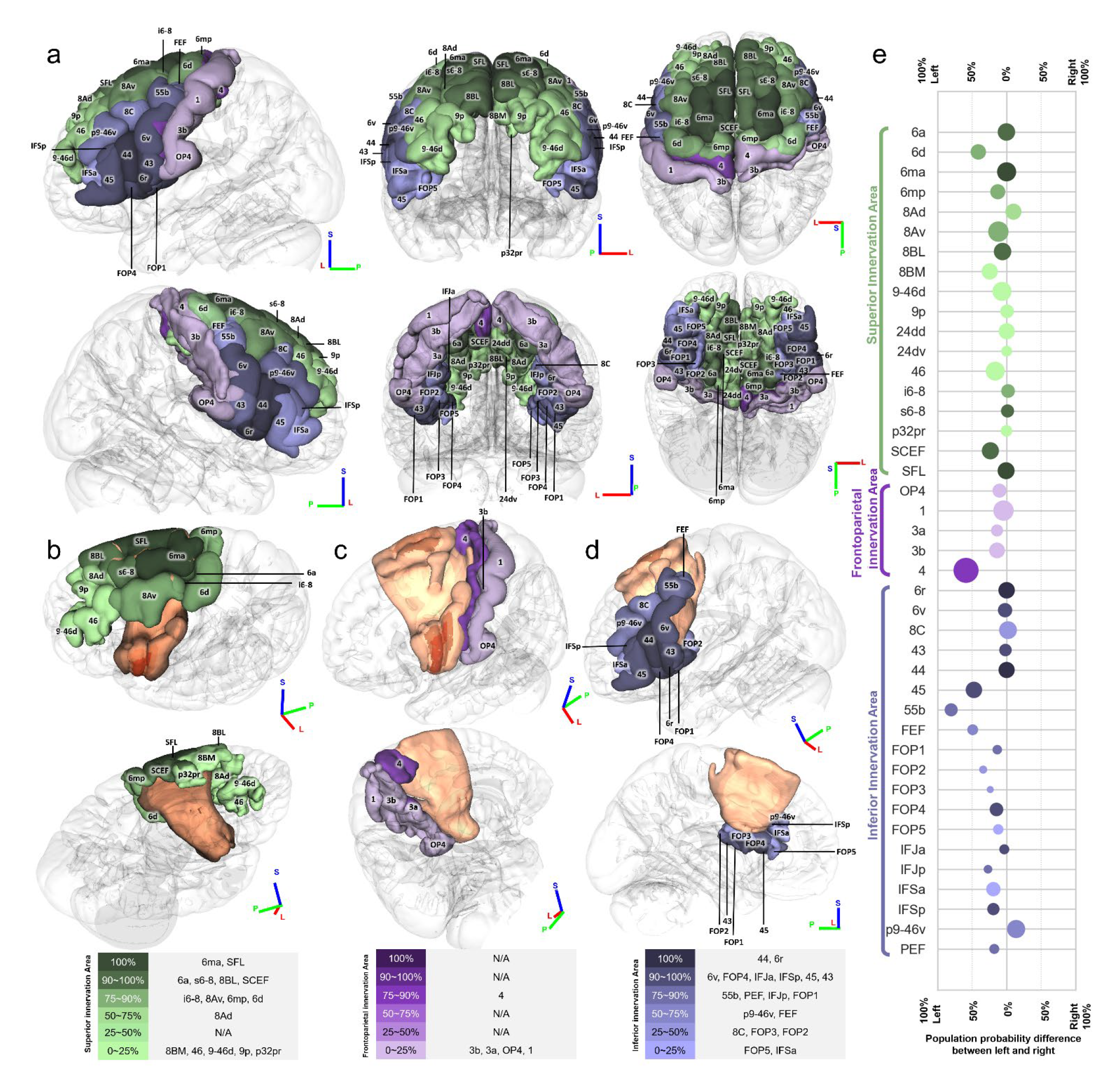
Population probabilities of the innervation of cortical regions and the FAT. (a) Green regions innervate the superior part of the FAT, while purple regions innervate the frontoparietal part, and blue regions innervate the inferior part. The darker color represents higher population probability. (b) The orange area is the FAT with the different shades representing Q1, Q2, and Q3. The FAT shows substantially connection with the areas in the superior frontal gyrus. (c) Other than area 4, other cortical regions have low population probabilities. (d) The areas with high population probabilities are mainly in the inferior frontal gyrus, including the pars opercularis and pars triangularis. (e) The population probability difference shows that the individual difference of most of the innervated areas are left biased. The surface area of the circle indicates the volume of the area.

### Tract-to-region connections

The findings pertaining to the connectivity of the FAT are presented in Fig. 5, a Sankey flow diagram that portrays the probabilities of the same source population, as previously demonstrated in Fig. 4. Fig. 5a and 5b illustrate the connections in the left and right hemispheres, respectively. The saturation of each region’s color in the diagram corresponds to its population probability. The colors used in Fig. 5 are also utilized to represent the superior, inferior, and frontoparietal innervation with the FAT, in green, blue, and purple, respectively. The diagram reveals that the inferior areas exhibit greater innervation with the FAT than the superior regions.

**Figure 5.**
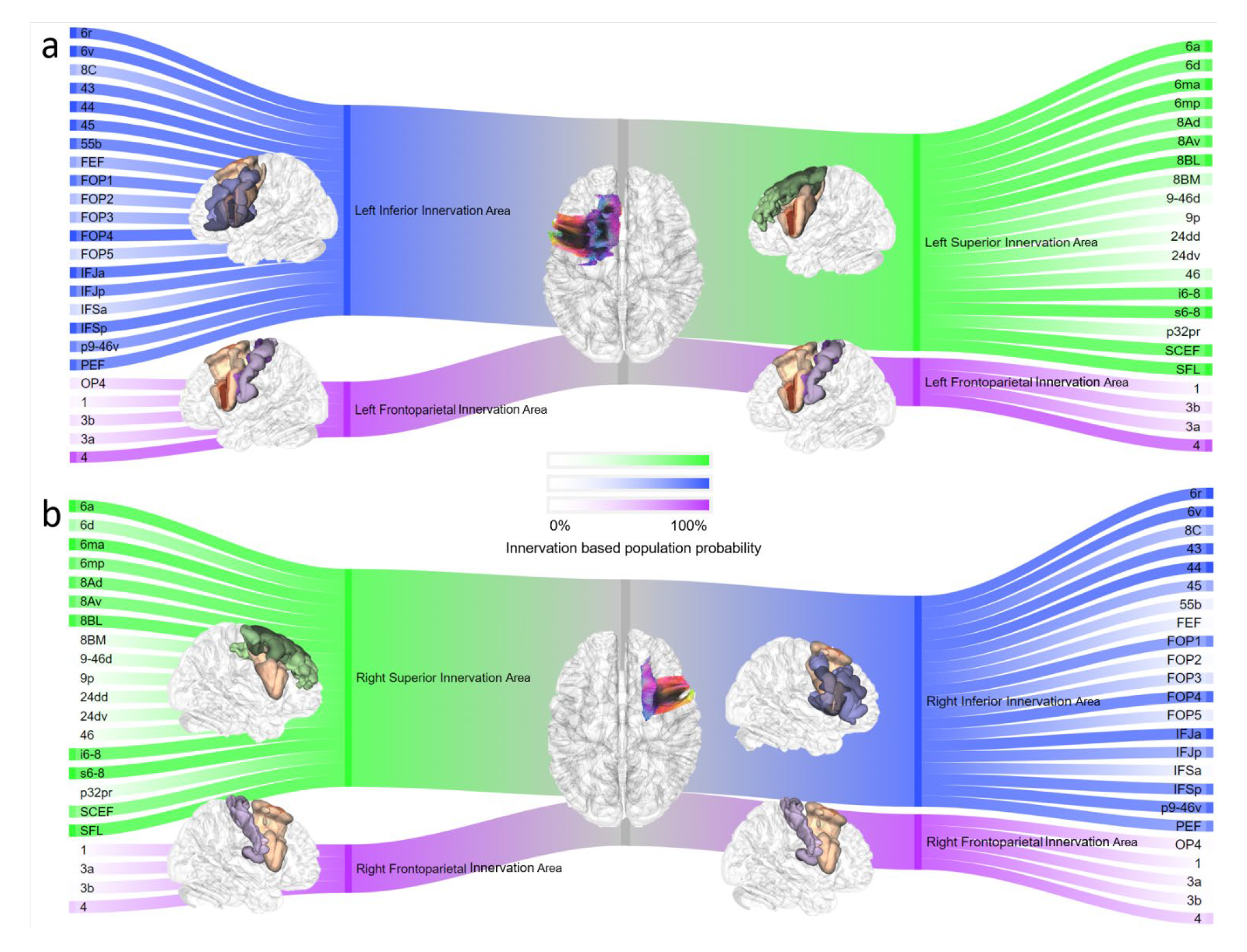
Sankey flow diagram showing the innervated cortical regions of the FAT. (a) The color saturation represents the population probability of cortical regions. This figure is showing the left FAT and the cortical regions it innervates. (b) This figure is showing the right FAT and the cortical regions it innervates.

### Structure-function correlation

In Fig. 6, we presented the relationship between the frontal aslant tract (FAT) and language-related functions, specifically the Oral Reading Recognition test and Picture Vocabulary test designed by the NIH toolbox. To examine the relationship, we used a linear mixed-effects model to regress participants’ covariates on the functions. Sex was treated as a random effect term, and other covariates as fix effect terms, as conducted in many language performance studies. Our results showed no significant correlation in the Oral Reading Recognition test. While not statistically significant, we also noted a near significant p-value in the inferior innervation area of the right FAT (p=0.066) in the Oral Reading Recognition test. The fractional anisotropy (FA) does not show correlation.

**Figure 6.**
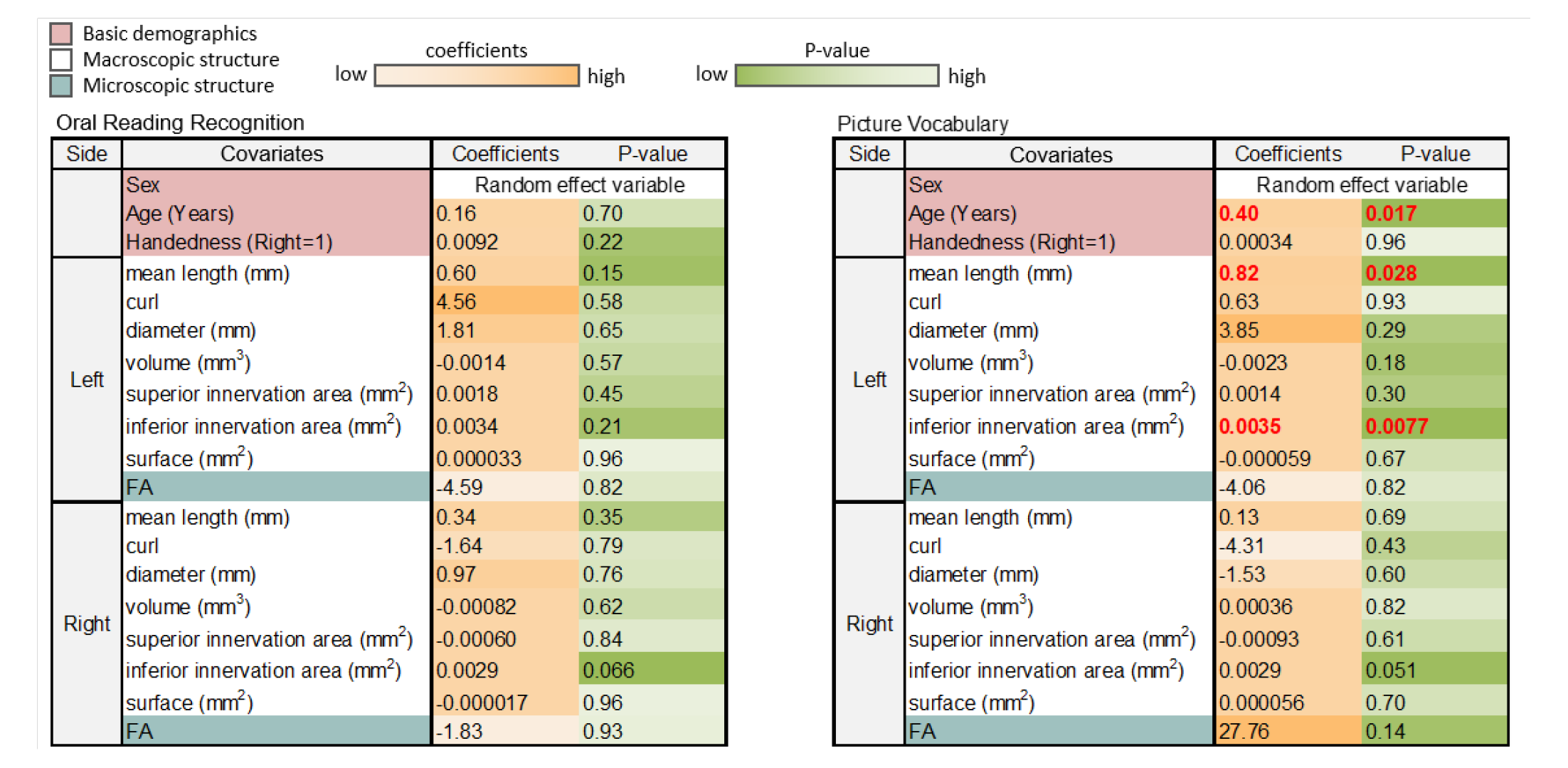
Results of the correlation analysis between the frontal aslant tract (FAT) and language-related functions. The Oral Reading Recognition test showed no significant correlation, while the Picture Vocabulary test exhibited a highly significant correlation with the size of the left FAT’s inferior innervation area (p=0.0077). Other correlations, including the mean length of the left FAT and FA values, were not statistically significant.

On the other hand, the Picture Vocabulary test showed a highly significant correlation with the size of left FAT’s inferior innervation area (p=0.0077). It is noteworthy that there is also a less significant correlation with the mean length of the left FAT (p=0.028), but likely owing to the confounding effect of the innervation area (a larger innervation area will lead to longer mean length). We observed that the inferior innervation area of the *right* FAT had a borderline p-value close to significance (p=0.051). There is also a lower p-value observed between FA of the right FAT and Picture Vocabulary (p=0.14), but the correlation is not statistically significant. Our overall result suggests that the function of FAT is related to language function examined by Picture Vocabulary test.

### Segmentation

Following our previous analysis, the FAT could be segmented into the anterior (blue) and posterior FAT (red) based on their respective connecting regions, as shown in Fig. 7a. The segmentation was based on the inferior frontal counterparts’ connecting regions, since the superior area of the both bundles connect with SFG, lacking an obvious segmentation. Fig. 7b anatomically depicts the curved shape of the FAT and its surrounding brain regions. Similar to the 3D model we created, the anterior FAT primarily innervates pars opercularis and partly innervates pars triangularis, while the posterior FAT innervates premotor cortex. Fig. 7c shows the white matter fiber pathways surrounding the FAT tracked using MRI data from an HCP young adult subject (#161731). The posterior FAT displayed more innervation with its surrounding white matter pathways comparing to the anterior FAT, particularly with the SLF II and SLF III. The CST, shown in purple, is located posterior and runs parallel to the FAT, while the AF, SLF II, SLF III run in the perpendicular direction and are more peripheral than the FAT.

**Figure 7.**
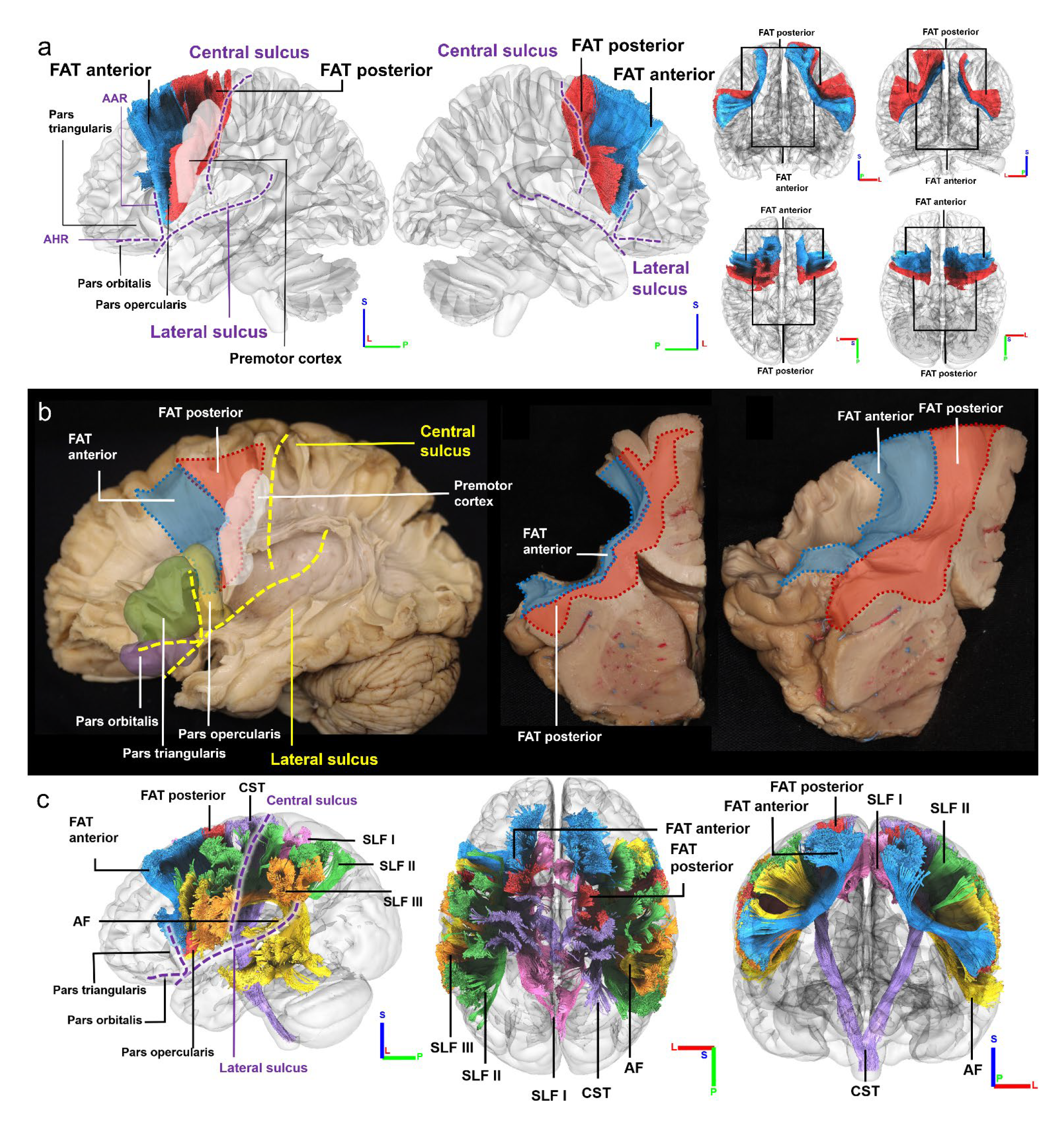
Segmentation of population-averaged FAT. (a) FAT can be segmented into the anterior FAT (blue) and posterior FAT (red). The posterior FAT innervates to the inferior frontal gyrus (IFG), specifically to the pars opercularis. (b) Posterior FAT innervates the premotor cortex (white), whereas the anterior FAT innervates the pars opercularis (yellow), pars triangularis (green), and their anterior structures. (c) Compared to the anterior FAT, the posterior FAT has more areas innervated with the surrounding pathways. The location of FAT runs more medial to other surrounding structures, such as SLF II (green), SLF III (red) and AF (yellow), whereas SLF I (pink) is medial to the FAT, and CST (purple) is positioned at the posterior side of the FAT.

## Discussion

In this study, we analyzed the individual differences of the frontal aslant tract (FAT) coverage in young adults using population probabilities and HCP-MMP atlas to define the cortical regions. The findings indicated that there are large individual differences in the anterior FAT in the superior and middle frontal gyrus, and the superior part of the FAT has larger individual differences compared to the inferior part. The left FAT has more coverage than the right, and the distinction between the left and right is greater in the superior and posterior side of the FAT coverage. The population probability of innervation with cortical regions was also analyzed, revealing that the size of inferior innervation areas is significantly correlated with Picture Vocabulary function. Furthermore, the left FAT inferior innervation areas have a significant correlation with Picture Vocabulary function, indicating the structural and functional importance of the left FAT in language processing. On the other hand, FA, representing the anisotropy of FAT, shows no significant correlation with the functioning of FAT.

### Individual differences

Our study is unique in that it is the first to analyze differences in FAT across a large cohort of 1065 subjects from the Human Connectome Project. White matter tracts are known to have individual differences, and these differences may be attributed to genetic or environmental factors ^11^. The structure of the FAT, which runs from the superior frontal gyrus to the inferior frontal gyrus, has been previously identified by Catani et al. ^3^. The original studies did not study its individual differences; however, our study found substantial individual differences in the FAT, which are not equally distributed. First, fewer individual differences are found in the inferior portion of the FAT. This could be related to the fact that FAT is funnel-shaped and has less dispersion of the white matter tracts in the inferior innervation area. Previous studies have revealed that the FAT innervate pars opercularis or pars triangularis ^3,6^. What we found is consistent: over 90% of the subjects had FAT connections in both pars opercularis and pars triangularis, indicating that most individuals have FAT connections in both areas. Furthermore, our study revealed additional information: the population-averaged FAT overlaps more with area pars opercularis than area pars triangularis. This finding has important implications for our understanding of language processing in the brain. The pars opercularis and pars triangularis are two regions of the frontal cortex that have been implicated in different aspects of language processing. An fMRI research revealed that the pars opercularis has been linked to syntactic processing, while the pars triangularis has been linked to semantic processing ^12^. The fact that the FAT overlaps more with the pars opercularis suggests that it may play a greater role in syntactic processing than semantic processing. This is consistent with previous research suggesting that the FAT is involved in the production and comprehension of semantic aspects of language ^1,13,14^. On the other hand, the fact that the overlap with the pars triangularis is lower suggests that the FAT may play a lesser role in syntactic processing. Understanding the specific neural circuits involved in different aspects of language processing may help to develop more targeted interventions for individuals with language disorders. Additionally, these findings may contribute to our understanding of the relationship between brain structure and function, and how individual differences in brain structure may relate to individual differences in language processing abilities.

Another key observation is our study is that the posterior FAT, which has less variability, connects the premotor cortex, whereas the anterior FAT, with greater variability, connects the inferior frontal gyrus. This is an interesting finding and has potential implications for our understanding of the functional organization of the brain. The premotor cortex is involved in planning and executing motor movements, while the inferior frontal gyrus is known to play a role in language processing and executive function ^15^. One possible explanation for the difference in variability between the anterior and posterior FAT is that the premotor cortex may be more genetically and developmentally constrained than the inferior frontal gyrus. This could mean that there is less room for individual variability in the connections between the premotor cortex and posterior FAT.

Another possible explanation is that the anterior and posterior FAT play different roles in the functional organization of the brain. For instance, the anterior FAT may be more involved in language processing and executive function as well as inhibitory control, while the posterior FAT may be more involved in motor planning. Overall, different mechanisms underlying the development and organization of different regions of the brain may have implications for our understanding of individual differences in brain structure and function. Further research is needed to better understand the functional implications of the variability in the connections between the FAT and different regions of the brain, and how this variability relates to individual differences in cognitive and motor function.

### Structural lateralization

In our study, our analysis showed significant left-lateralization of the FAT. Our population analysis showed that the left FAT is in average larger than the right side, and it also has larger individual differences, specifically at the inferior part of the FAT. The structural lateralization has implications for our understanding of brain asymmetry and its relationship to cognitive function. Asymmetry in brain structure and function has been recognized as an important aspect of human cognition ^16^, and understanding the neural mechanisms underlying this asymmetry can help us better understand how the brain works. The fact that the left FAT is larger and has more individual differences, particularly in the inferior part, may suggest that it is more involved in language-related processes, such as semantic processing or word retrieval, than the right FAT. However, it is important to note that the superior portion of the FAT, which stems from the pre-SMA and SMA, did not show as much left-right difference as the inferior portion. This suggests that the role of the superior portion of the FAT may not be as strongly lateralized as the inferior portion. Further research is needed to determine the specific functional roles of the superior and inferior portions of the FAT, and how these roles relate to lateralization and individual differences.

### Structure-function correlation

One explanation for the structural lateralization of the FAT is that it reflects functional specialization in the brain. Previous research has shown that the FAT in the left hemisphere is more involved in language processing, while in the right hemisphere it is more involved in inhibitory control ^1,2^. Our structure-function analysis also showed that the inferior innervation area of the left FAT is significantly correlated with the performance of the Picture Vocabulary test. This result is consistent with previous studies that have shown that the left FAT is associated with initiating speech and language fluency, while the right FAT is involved in supporting inhibitory control ^1,2^. In overall, the left FAT may play a particularly important role in language processing, while the right FAT may be more involved in cognitive control processes. Interestingly, another study showed that the right laterality in the FAT was demonstrated with connections of the pars opercularis to the pre-SMA in 5-8 year-old children ^17^. This suggests that the functional lateralization of the FAT may change over the course of development, with the right side becoming more involved in certain cognitive processes as children mature.

### FAT in dual stream models

Our comprehensive structure-function analysis revealed compelling correlations between FAT morphology and language function, providing robust support for the role of FAT in language processing. However, when examining two specific language functions, namely the Oral Reading Recognition and Picture Vocabulary tests, intriguing distinctions emerged. Only the Picture Vocabulary test demonstrated a highly significant correlation with the surface area of the left FAT’s inferior innervation region. Additionally, the right FAT’s inferior innervation area exhibited a borderline significance. These findings prompt a deeper exploration into the intricate dynamics between FAT morphology and language tasks.

To shed light on these results, we turn to the influential dual stream model proposed by Hickok and Poeppel ^18^. According to this model, speech processing involves two distinct neural pathways: the ventral stream, responsible for speech comprehension, and the dorsal stream, facilitating the transmission of signals to articulatory networks. Delving into the specifics, the Picture Vocabulary test predominantly engages the ventral stream by requiring individuals to recognize and associate visual representations with their corresponding verbal labels or meanings. Consequently, the significant correlation observed between the surface area of the left FAT’s inferior innervation region and the Picture Vocabulary test suggests that the connectivity of FAT within the ventral stream contributes significantly to tasks reliant on comprehension processing. While the study conducted by Saur, Kreher et al. ^19^, primarily examined activations within the two streams using functional MRI (fMRI) and traced white matter fibers associated with them using diffusion tensor imaging (DTI), the specific involvement of FAT in the ventral stream was not explicitly mentioned. However, given its close neuroanatomical proximity, it is highly plausible that FAT plays a vital role within the ventral stream, aligning with its function in semantic processing tasks like the Picture Vocabulary test. On the other hand, the dorsal stream projects towards the inferior parietal and posterior frontal lobes through the arcuate fasciculus (AL) and superior longitudinal fasciculus (SLF), predominantly engaging in phonological and orthographic processing. The lack of a significant correlation between FAT morphology and the Oral Reading Recognition test suggests a less pronounced association of FAT with the dorsal stream.

The contrasting correlations between FAT and different language tasks can be elucidated by considering the specialization of the ventral and dorsal streams within the dual stream model. The prominent connectivity of FAT within the ventral stream likely contributes to tasks involving semantic processing, such as the Picture Vocabulary test. In contrast, the influence of FAT on reading tasks may be comparatively diminished due to the dominant involvement of the dorsal stream in those specific processes. These nuanced findings underscore the complexity of FAT’s role in language processing and emphasize the importance of considering distinct neural pathways when studying its functional implications.

### Role in neurosurgery

Our study has significant implications for the field of neurosurgery, particularly in the context of low-grade gliomas located in the superior frontal gyrus, including the supplementary motor area (SMA) and pre-supplementary motor area (preSMA). These types of tumors are commonly found in young patients, underscoring the importance of considering the frontal aslant tract (FAT) during presurgical planning and intraoperative procedures when removing brain lesions in this area. A thorough understanding of the FAT’s fiber predominance, orientation, and volume is crucial in the intraoperative setting to ensure a safe surgical approach and minimize the potential for unnecessary damage to the brain, thereby reducing the risk of long-term complications.

Previous research has shed light on the significance of the left FAT in various conditions. For instance, a study employing direct axonal stimulation and postoperative tractography investigated the role of white matter tracts, specifically the left FAT, in stuttering ^20^. The findings revealed that the left frontal aslant tract plays a pivotal role in stuttering and may be involved in a cortico-subcortical circuit responsible for speech motor control. Another study reported that a unilateral lesion encompassing the insular and the FAT resulted in bilateral paralysis of the facial-lip-pharyngeal-laryngeal musculature ^5^. Additionally, disruption of the fronto-parietal and frontal cortico-subcortical connectivity, including the FAT, was found to be correlated with long-lasting impairments of executive functions. These insights are invaluable for surgical planning and predicting neuropsychological disorders in brain tumor surgery ^21^.

There is a general consensus among researchers that the left FAT is intimately involved in speech initiation. This aligns with the role of the posterior segment of the FAT, which connects the SMA and preSMA to the facial-lip motor cortex, contributing to the initiation of speech. However, it is worth noting that our study did not specifically examine language initiation due to the specific language tests employed. Nonetheless, our correlational evidence suggests that

FAT may play a role in the semantic/ventral stream of language processing. This seems to match the role of the anterior segment of FAT that connects the superior frontal lobe to the inferior frontal regions, including Broca’s area, which is known for its involvement in language production and comprehension. Further research is warranted to explore and elucidate the distinct functions of the anterior and posterior segments of the FAT more comprehensively.

Importantly, our study revealed substantial individual differences in the left anterior FAT, highlighting the potential risk for language deficits following surgery in this region. Therefore, for lesions located in the left inferior frontal lobe, employing an individualized mapping of the FAT could prove beneficial in reducing post-surgical functional deficits and optimizing patient outcomes.

### Limitation

However, several limitations still exist in our study. We cannot conduct a population-based study using cadavers. The population studies only used imaging data from the human connectome project. Furthermore, tractography has errors, and the indistinct boundaries between white matter pathways in specific individuals would lead to an inferior distinction. False continuations could occur with SLF and other nearby pathways. Lastly, we haven’t conducted a functional mapping in this study to examine our functional claim about anterior and posterior FAT. This could be done by fMRI or intraoperative functional mapping.

Although language-related functions have been analyzed in this study, motor planning functions are still to be analyzed in the future. Language fluency is said to be associated with the FAT in former studies ^1,2^, which is also not performed in this study and needs further analyzation. For future clinical applications, the FAT segmentation could inform possible eloquent area, especially the anterior FAT.

In addition to examining individual differences in the topological distribution of the FAT, future studies could focus on investigating how these differences develop over time. Research could explore whether there are developmental differences in the FAT between children and adults, as well as the environmental and genetic factors that may contribute to these differences. Further studies could also investigate how the FAT varies in individuals with neurodevelopmental disorders, such as autism or schizophrenia. Another potential area of investigation is the functional significance of the variations in the FAT, including how they relate to individual differences in cognitive and behavioral functions. Finally, studies could explore how the variations in the FAT relate to the structural and functional connectivity of other neural networks in the brain, and how these networks are affected in neurological and psychiatric disorders.

## Acknowledgements

The authors were supported by NIH grant R01 NS120954, R56 MH113634, and R01 DC013803. Data were provided in part by the Human Connectome Project, WU-Minn Consortium (Principal Investigators: David Van Essen and Kamil Ugurbil; 1U54MH091657) funded by the 16 NIH Institutes and Centers that support the NIH Blueprint for Neuroscience Research; and by the McDonnell Center for Systems Neuroscience at Washington University.

During the preparation of this work the authors used ChatGPT 3.5 (OpenAI) to revise the manuscript. After using this tool/service, the authors reviewed and edited the content as needed and take(s) full responsibility for the content of the publication.

## Author Contributions

WJL and JBM performed the analyses and contributed to the writing of the manuscript. DFC, TJ, MN, RG, and YA conducted the cadaver dissection and provided critical review of the manuscript. JFM initiated the study, reviewed the manuscript, and provided guidance. FCY contributed to the writing of the manuscript and oversaw the study.

## Materials and Methods

### MRI Acquisition

We utilized a dataset consisting of 1065 subjects, which was obtained from the Human Connectome Project and made available by the WashU consortium ^22^. The diffusion data were acquired using a multishell scheme, incorporating three b-values (1000, 2000, and 3000 s/mm^2^), with 90 directions in each shell. The spatial resolution of the acquired images was 1.25 mm isotropic. The ac-pc line of the ICBM152 space is the standard of alignment. The diffusion data and b-table were linearly rotated and interpolated with cubic spline interpolation at 1mm. Q-sampling imaging with a diffusion sampling length ratio of 1.7 is used to reconstruct the rotated data, which were later used in automated tractography ^23^. To ensure the accuracy and orientation of the b-table, an automatic quality control routine ^24^ was employed.

### Automated tractography

The automated tractography pipeline in DSI studio (http://dsi-studio.labsolver.org) is used to map the 52 white matter bundles of each subject. This pipeline is a combination of deterministic fiber tracking algorithm ^25^, topology-informed pruning ^26^, and randomized parameter saturation ^9^, with trajectory-based tract recognition ^9^ being an integrated interface, as detailed in a recent study ^8^. Following the mapping process, the white matter bundles of the 1065 subjects were exported to the ICBM152 2009 nonlinear space to examine the population variation of the FAT. The analysis was conducted in DSI Studio package, which is publicly available at http://dsi-studio.labsolver.org.

### Dissections

The human brain of 5 neurological healthy subjects were studied and compared with the results of the fiber tracking done by not only the manual processing but the automatic processing as well. Dissections were performed by expert neurosurgeons (TJ, MN, RG, and YA). Human brains were fixed in 10% formalin for 28 days and frozen for 14 days. Dissections were carefully performed from superficial to deep, starting at the grey matter until white matter fibers were encountered. The anatomical dissections of the FAT were placed in a high degree of difficulty. This is because the fibers of this tract are highly anatomically related to the fibers of the SLF tract, which crosses the bundle and merges in a complex way. The separation of the fibers was done in a way to preserve the integrity of the FAT without destroying the underlying or overlapping tissue.

### Tract-to-region connectome

The tract-to-region connectome, based on the HCP-MMP parcellations from a previous study ^8^, was utilized in our analysis. The HCP-MMP in ICBM152 space was obtained from an asymmetrical and improved reconstruction version of MMP 1.0 MNI projections from NeuroVault (https://identifiers.org/neurovault.collection:1549). The file is further edited with DSI Studio and shared on DSI Studio website: http://dsi-studio.labsolver.org. By mapping the intersection between the white matter bundles and cortical regions at the voxel level, a binary tract-to-region connection matrix was generated. The aggregation of binary matrices across the 1065 subjects results for the population probability of the tract-to-region connection, which is the tract-to-region connectome (available at http://brain.labsolver.org).

